# DICAR-JP Regulates Ribosome Migration, a New Theory of Mitochondrial Protein Production

**DOI:** 10.1101/2025.04.11.648480

**Authors:** Jun Zhang, Xueting Yu, Sihui Zheng, Gan Qiao, Chao Zhang, Siyi Tang, Xiaoping Gao, Yu Wang, Yajun Yu, Jun Cheng, Ming Lei, Pengyun Li, Yan Yang, Chunxiang Zhang, Qiong Yuan

## Abstract

**Background:** DICAR plays a cardioprotection of diabetic models. DICAR-JP, a short sequence of DICAR, may represent DICAR’s active functional domain. NACα is necessary for the heart tissue development. Addtionally, OGDHL is a metabolism regulator. In this study, we examine a new mechanism of DICAR via NACα/OGDHL on regulating cardiomyocyte metabolism in DCM. We also sought to elucidate the function of DICAR/DICAR-JP in modulating NACα-dependent production of OGDHL nascent peptides and their mitochondrial translocation, and we also aimed to optimize the DICAR-JP sequence to be better therapy on DCM.

**Methods:** SPR was to investigate the binding interaction between DICAR-JP and NACα. The function of DICAR-JP and DICAR-JP45 on OGDHL nascent peptide transfection was detected by in vitro translation approach. Biotin-DICAR-JP was transfected to study the function of DICAR-JP, NACα and OGDHL. Untargeted metabolomics was utilized to characterize the metabolic reprogramming of cardiomyocytes. AAV9-DICAR-JP and DICAR-JP45 were constructed and administered to db/db mice for 1-2 months to assess their cardioprotective effects detected by Echocardiography confirmed mouse cardiac function.

**Results:** Our findings demonstrated that DICAR-JP be the functional domain that interacts with NACα to regulate OGDHL nascent peptide expression. Disruption of OGDHL expression reversed the metabolic reprogramming observed in diabetic cardiomyocytes, highlighting its crucial role in maintaining cardiac metabolic homeostasis. DICAR-JP facilitated the translocation of OGDHL nascent peptides from the cytoplasm to the mitochondria, leading us to hypothesize that DICAR-JP plays a key role in regulating ribosomal migration from the endoplasmic reticulum to the mitochondria. We refer to this process as the ‘Ribosome Migration’. In our studies using AAV9-mediated DICAR-JP and DICAR-JP45 overexpression in heart tissue. DICAR-JP and DICAR-JP45 both exhibited significant cardioprotective effects against diabetic cardiomyopathy (DCM), comparable to those observed with Empagliflozin.

**Conclusions:** In a conclusion, we propose a new endogenous nucleic acid candidate drug library, and a new molecular framework, the ‘Ribosome Migration’, which implicates the DICAR-JP/NACα/OGDHL nascent peptide axis in metabolic reprogramming related to DCM. This theory constructs a new mitochondrion protein from nuclear original protein. Moreover, DICAR-JP45 shows strong potential as a nucleic acid-based therapeutic candidate for treating DCM.

**Novelty and Significance:** *What Is Known?:* DICAR is a new protective circular noncoding RNA for diabetic cardiomyopathy. NACα is located in ribosome and regulates new peptide production, which is also a key factor in heart tissue development. OGDHL can regulate different energy metabolism in mitochondria.

*What New Information Does This Article Contribute?:* DICAR-JP is the functional domain that interacts with NACα to regulate OGDHL nascent peptide expression. There is an interaction between DICAR-JP and NACα. DICAR-JP sequence is protected by NACα and NACα function is mediated by DICAR-JP. We called this ‘RNA functional domain’. OGDHL expression reversed the metabolic reprogramming observed in diabetic cardiomyocytes, highlighting its crucial role in maintaining cardiac metabolic homeostasis. DICAR-JP takes part in tanscription and translocation of OGDHL nascent peptides from the cytoplasm to the mitochondria via plasma ribosome. We call this process the ‘Ribosome Migration Theory’. DICAR-JP and DICAR-JP45 both exhibited significant cardioprotective effects against diabetic cardiomyopathy (DCM), comparable to those observed with Dapagliflozin (DAPA). Our study revealed a novel molecular framework, the ‘Ribosome Migration Theory’, which implicates the DICAR-JP/NACα/OGDHL nascent peptide axis in metabolic reprogramming related to DCM. Moreover, DICAR-JP45 shows strong potential as a nucleic acid-based therapeutic candidate for treating DCM.

---

Diabetic cardiomyopathy (DCM) is a significant and often overlooked complication of diabetes, contributing substantially to the burden of cardiovascular disease.^1,2^ Clinical trial results indicate that the mortality and morbidity rates among patients with heart diseases who also have diabetes are three times higher than those without diabetes. This increased risk can be attributed to various factors associated with diabetes, including insulin resistance, chronic inflammation, and the accumulation of advanced glycation end products (AGEs), all of which lead to myocardial injury and dysfunction.^3^ Recent research has demonstrated the protective effects of Dapagliflozin (Dapa),^4^ a sodium-glucose co-transporter-2 (SGLT2) inhibitor, particularly in patients with heart failure with preserved ejection fraction (HFpEF).^5^ Given the intricate relationship between diabetes and cardiovascular health, advancing effective therapies for DCM is a critical area of research. By uncovering the pathophysiological mechanisms driving DCM, scientists aim to develop targeted treatments that not only improve clinical outcomes but also enhance the quality of life for patients affected by this debilitating condition.

DICAR, also known as circTULP4, is a newly identified circular RNA (circRNA) that has shown a beneficial role in mitigating the effects of diabetes cardiomyopathy (DCM) in a mouse model.^6^ Research indicates that DICAR is significantly downregulated in pancreatic islets^7^ and neurons,^8^ suggesting that its expression may be linked to the metabolic and neurodegenerative complications associated with diabetes. The primary mechanism by which DICAR exerts its protective effects in DCM is through its interaction with valosin-containing protein (VCP).^6^ This crucial binding event regulates the degradation of Med12, a key component of the Mediator complex involved in nuclear transcription. The degradation occurs via ubiquitination processes in the endoplasmic reticulum (ER),^9^ illustrating how DICAR may influence gene expression and cellular stress responses. In our previous research, we further elucidated the role of DICAR-JP, a functional part of DICAR that covers the junction site of the circRNA. We found that DICAR-JP plays a critical role in the protective function of DICAR, particularly regarding its potential to inhibit pyroptosis—a form of programmed cell death characterized by the release of inflammatory cytokines—in cardiomyocytes impaired by AGEs.^6^ Thus, both DICAR and DICAR-JP may be pivotal in inhibiting the onset and progression of DCM, providing a promising avenue for therapeutic intervention. Understanding the specific molecular pathways and interactions could lead to the development of novel strategies aimed at targeting circRNA pathways to enhance cardiac health in diabetic patients. Future research can focus on the clinical implications of these findings, exploring how modulation of DICAR levels can be translated into therapeutic benefits for individuals suffering from diabetes-related cardiac complications.

In our previous research, we discovered that DICAR binds to the NACα protein, a critical component of the nascent polypeptide-associated complex (NAC). This complex is essential for interacting with newly synthesized proteins as they emerge from the ribosomal tunnel exit.^10,11^ NACα plays a vital role in ensuring the correct targeting of these nascent polypeptides, as it competes with the signal recognition particle (SRP) to prevent the mistargeting of cytosolic and mitochondrial proteins to the ER.^12,13^ The precise localization of NACα at the ribosomal tunnel exit underscores its regulatory function in peptide production^14^ and in mediating the translocation of these peptides into mitochondria, where they are crucial for various metabolic processes.^15^ Moreover, NACα has been shown to interact with the E3 SUMO ligase, mammalian Mms21/Nse2. Knockdown of Nse2 expression has been linked to significant impairments in myogenic differentiation, which is crucial for the proper development of muscle tissue.^16^ This inhibition is accompanied by a partial blockade of the nuclear-to-cytoplasmic translocation of the skNAC-Smyd1 complex,^16–18^ leading to the retention of this complex within promyelocytic leukemia (PML)-like nuclear bodies. Such retention disrupts sarcomerogenesis, the process by which muscle fibers are formed and organized, indicating that the nuclear and cytosolic roles of the NACα-Smyd1 complex are finely balanced and may be regulated through sumoylation, a post-translational modification that affects protein interactions and stability. Furthermore, studies utilizing a Drosophila model have demonstrated that the knockdown of NACα disrupts cardiac developmental remodeling, ultimately resulting in the absence of a functional heart.^12^ This finding highlights the critical role of NACα in cardiac development and suggests that proper regulation of its activity is essential for maintaining normal cardiac morphology and function. Understanding the associated mechanisms of DICAR and NACα may provide valuable insights into potential therapeutic targets for diseases associated with impaired cardiac development and function.

In the same CHIRP-MS data, we also found that DICAR can bind with oxoglutarate dehydrogenase (OGDHL), a glycosyltransferase.^19,20^ This protein modulates glutamine metabolism to influence tumor progression.^20^ OGDHL is a nuclear original protein and located in the mitochondrion,^21,22^ and is also found to induce DNA impairment.^23^ Recently, the M2 AChR/OGDHL/ROS axis was found to play roles in inhibiting the pyroptosis of heart injured by myocardial ischemia-reperfusion.^24^ Therefore, in this study, we would like to confirm that DICAR-JP promotes OGDHL mediating energy reprogramming in DCM.

Therefore, in this study, we aimed to explore the function on DICAR/DICAR-JP expression; DICAR/DICAR-JP’s regulation on NACα which has effects on OGDHL transcription, translocation and mitochondria energy reprogramming.

## Methods

### Data Availability

All supporting data of this study are available from the corresponding author upon reasonable request. For detailed experimental methods, materials, and statistical analyses, please see the Supplemental Material and Major Resource Table.

The following citations are for the expanded Materials and Methods and Major Resource Table sections.

### Animal care and experimental procedures

C57BL/KsJ wild-type (WT) mice, aged 8 weeks, and C57BL/KsJ leptin receptor-deficient (Leprdb/db) mice of both sexes, exhibiting blood glucose levels between 23 and 28 mM, were procured from the breeding colonies of GenePharmatech Company, which is originally affiliated with The Jackson Laboratory. DICAR^+/ −^ mice were generated at the Model Animal Research Center of Nanjing University, China (refer to the Results section for details). All animal experiments were conducted using age– and sex-matched controls. The mice were housed in a temperature-controlled environment (22 – 25 °C) with a 12-hour light/dark cycle, ensuring ad libitum access to food and water at the Animal Center of Wuhan University of Science and Technology.

### Generation and administration of AAV9

For detailed information, please refer to the Methods section in the Supplemental Material.

### Experimental protocol

All mice administered AAV9 were euthanized 8 weeks following viral injection, at 17 weeks of age, to assess the long-term effects of the viral transduction on cardiac function and overall physiology. Comprehensive evaluations were conducted both prior to and 8 weeks post-AAV9 administration to ensure a thorough understanding of the physiological changes induced by the treatment. These assessments included echocardiography to evaluate cardiac structure and function. Exercise tolerance tests were also performed to determine the impact of AAV9 treatment on the physical capabilities of the mice, thus providing insights into their overall cardiovascular fitness. These metrics are critical for understanding the overall health status of the animals and for correlating structural changes in the heart with systemic physiological effects. Following euthanasia, cardiac tissues were harvested and processed for various analyses. Furthermore, mass spectrometry (MS) analysis was utilized to profile protein expression and metabolomics within the cardiac tissue or, facilitating a deeper understanding of the molecular mechanisms underlying the observed physiological changes.

### Cardiomyocyte culture

For detailed information, please refer to the Methods section in the Supplemental Material.

### Western blotting

For detailed information, please refer to the Methods section in the Supplemental Material.

### Histological analysis

For detailed information, please refer to the Methods section in the Supplemental Material.

### Chromatin isolation by RNA purification–MS (CHIRP-MS)

For detailed information, please refer to the Methods section in the Supplemental Material.

### RNA Immunoprecipitation (RIP) Assay

For detailed information, please refer to the Methods section in the Supplemental Material.

### DICAR-JP sequence mutation

For detailed information, please refer to the Methods section in the Supplemental Material.

### Structured illumination microscopy (SIM)

For detailed information, please refer to the Methods section in the Supplemental Material.

### Untarget Metabolome analysis

For detailed information, please refer to the Methods section in the Supplemental Material.

### Statistical analyses

All data are presented as means ± SEM. Statistical analysis for the comparison of two groups was performed using a two-tailed unpaired Student’s t-test. To compare more than two groups, a one-way analysis of variance (ANOVA), followed by Tukey’s post hoc test was performed. Adjusted two-sided P-values were calculated, and P < 0.05 was considered to indicate statistical significance. All statistical analyses were performed with the GraphPad Prism Version 6 (GraphPad Software Inc., SanDiego, CA, USA) and SPSS package (SPSS Inc., Chicago, IL, USA).

## Results

### NACα is the target protein of DICAR in diabetic cardiomyopathy

In our recent study, we employed RIP followed by CHIRP-MS, and analyzed data using the KEGG pathway analysis. The primary focus was to understand the biological functions of DICAR, particularly its involvement in key metabolic pathways such as carbon metabolism, oxidative phosphorylation, and the tricarboxylic acid (TCA) cycle (Figure 1A). Our results strongly suggest that DICAR plays a pivotal role in regulating metabolic processes, including both protein and energy metabolism (Figure 1A). These findings emphasized DICAR’s potential as a metabolic regulator in cellular bioenergetics. In this context, our data also indicated a significant interaction between NACα and DICAR, particularly in the diabetic (db/db) mouse model when compared to the WT group (Figure 1B). Additionally, we used NACα antibody to confirm the interaction of DICAR and NACα, as the results showed in Figure 1C, NACα binded DICAR significantly. Based on these findings, we hypothesize that NACα may play a crucial role in the pathophysiology of diabetic cardiomyopathy (DCM), acting through its interaction with DICAR and affecting metabolic regulation. To further investigate this interaction, we employed RNAfold 2 to predict the secondary structure of DICAR (Figure 3D). Previous studies have identified the junction region of DICAR (DICAR-JP) as a critical functional domain responsible for its binding with VCP for the ub-Protein degradation system. Using bioinformatics tools, we modeled the binding interaction between DICAR-JP and NACα, identifying four key amino acid residues—Lys-65, Glu-69, Ala-59, and Arg-98-at the binding interface (Figure 3E). In addition, we also mimic the binding model of DICAR-JP and NACα. In the model, we considerated that DICAR-JP bind with NACα and located at the side of ribosome tunnel exit (Figure 1F). These results confirmed that DICAR physically interacts with NACα and that the interaction is mediated through these specific binding sites. Collectively, these findings offered valuable insights into the molecular mechanisms of DICAR and NACα interactions, and suggested NACα’s potential as a critical player in DICAR-mediated metabolic regulation, particularly in pathological states such as diabetic cardiomyopathy.

**Figure 1.**
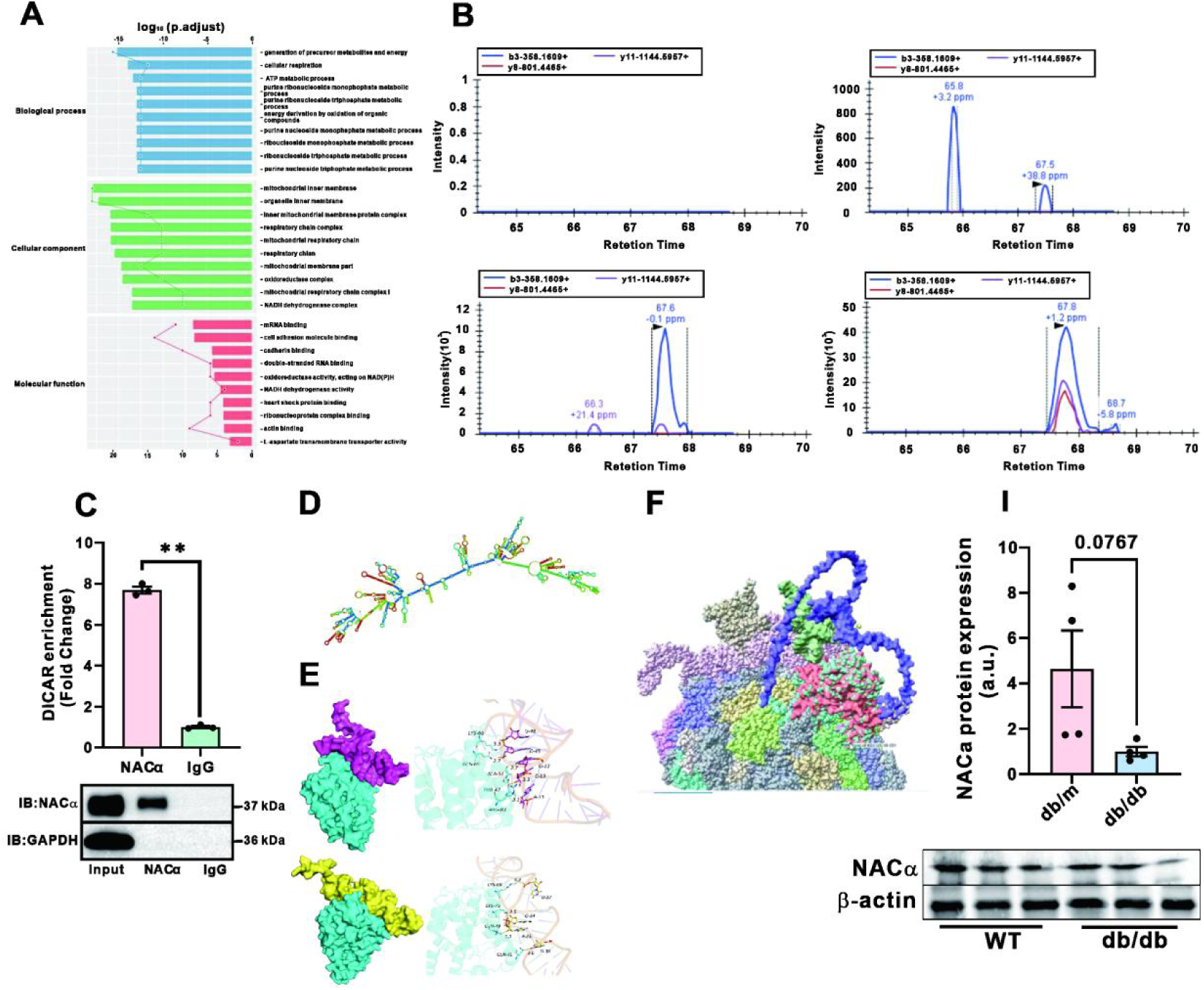
The interaction of DICAR/DICAR-JP and NACα. A: KEGG analysis of DICAR CHIRP-MS; B: Representative images of the NACα protein MS/MS spectrum detected by CHIRP-MS; C: RIP was used to identified the binding ability of NACα protein and DICAR in db/db mouse; D: RNAfold prediction of the secondary structure of DICAR; E: Bioinformatics used to mimic the binding modle of NACα and DICAR-JP; F: PyMOL bio-software prediction of the structure of DICAR-JP, NACα binding located in the ribosome; G: WB was used to detect NACα protein expressed in db/db mouse. N =3.

### The key sequence of DICAR on regulating NACα mediated OGDHL nascent peptide in ribosome

Given NACα’s known role in protein folding and its localization within the ribosome, we further studied the functional implications of this interaction by constructing a structural model of NACα and DICAR-JP within the ribosomal complex. In our CHIRP-MS analysis, we found that DICAR interact with the OGDHL protein (Figure 2A). Therefore, we transfected DICAR and DICAR-JP into AC16 to identified the function on combination of NACα and SRP54 proteins by Co-IP. However, we found that both of DICAR and DICAR-JP having no function on this complex formation (Figure 2B). In order to identified the function of NACα and DICAR-JP on OGDHL expression, we transfected NACα-OE, NACα-siRNA and DICAR-JP into AC16 to explore the total OGDHL expression. As the results showed in Figure 2C, OGDHL was downregulated in AGEs treated group, while NACα-OE reveresed the function of AGEs. Addtionally, NACα-siRNA inhibited OGDHL expression. However, DICAR-JP did not inhibite the function of NACα on OGDHL. Therefore, we considerated that DICAR-JP not play function on OGDHL protein expression without NACα. Additionally, we wonder to know that NACα and DICAR-JP on location of OGDHL expression, we separated the protein of mitochondrium, cell plasima and total protein. The results showed that OGDHL protein downregulated in total protein (Figure 2D) and mitochondirum (Figure 2E), however, in the plasmic protein, OGDHL did not changed (Figure 2F). To further verify DICAR’s impact on OGDHL expression, we utilized a cell-free protein transduction system to confirm the tranfect effect of DICAR-JP on OGDHL. As the results shown in Figure 2G, exgenouse DICAR-JP enhanced OGDHL protein transfection. Additionally, we transfected bio-DICAR-JP into AC16 and then collected the cell lysis to confirm the interaction of DICAR-JP and OGDHL. The results showed that DICAR-JP can significantly binds with OGDHL (Figure 2H). We cotransfected OGDHL, NACα and DICAR-JP into 293T cell. The results showed that OGDHL, DICAR-JP coexpression in the membrane of mitochondrion. NACα expressed location was not specific near mitochondrium, but all of the cell. Therefore, we constructed NACα plasmic including targeted mitochondrium. It was observed that NACα promote DICAR-JP function on OGDHL nascent peptides at the mitochondrial membrane and their translocation into the mitochondria. This mechanism of ‘simultaneous translation and translocation’ represents a novel new pathway for nuclear-encoded gene expression within mitochondria.

**Figure 2.**
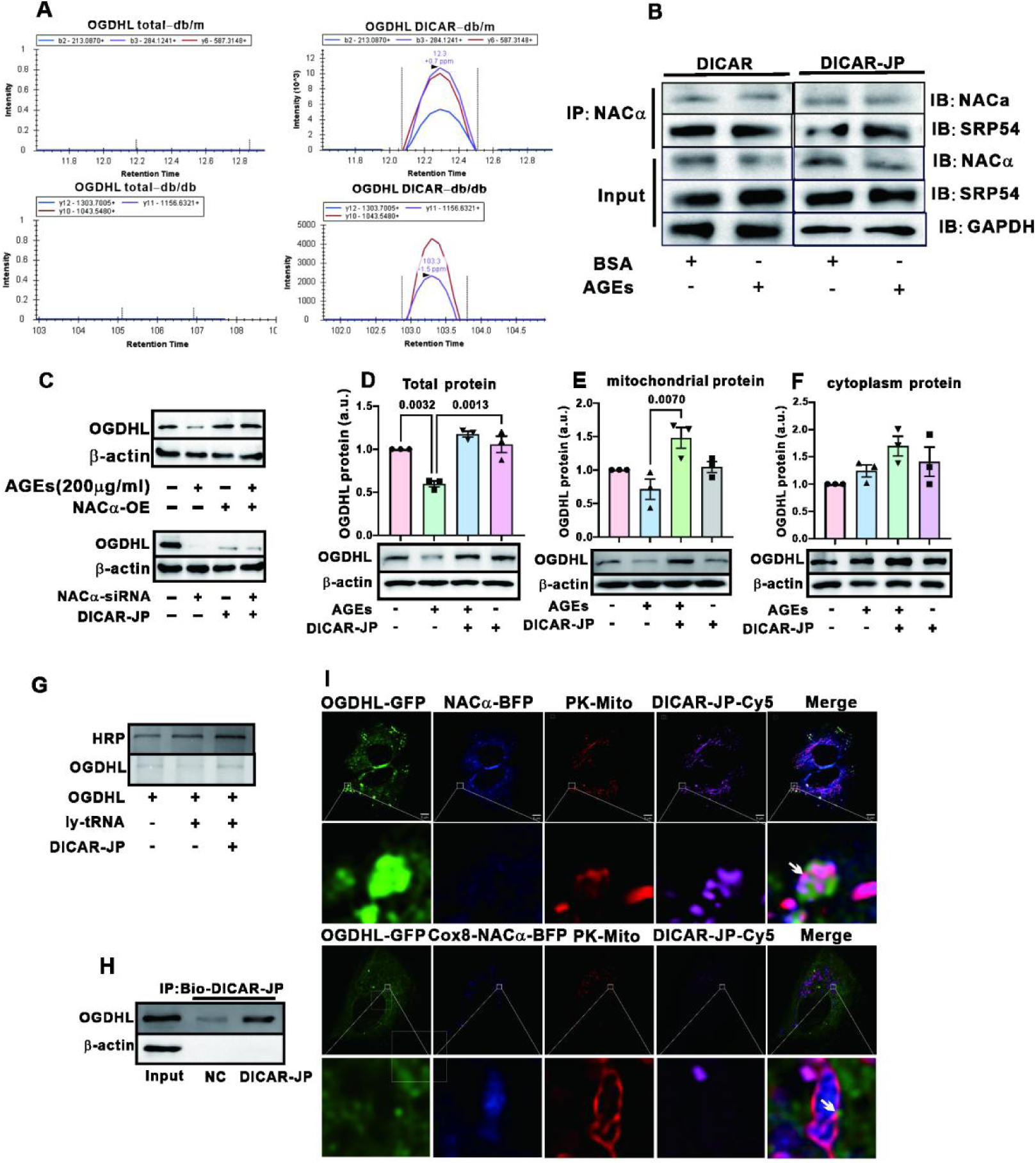
Effect of DICAR-JP on OGDHL nascent peptide production. A: Representative images of the NACα protein detected by CHIRP/MS spectrum; B: Co-IP was used to detect DICAR and DICAR-JP on combination of NACα and SRP54; C: OGDHL protein expression regulated by NACα-OE, NACα-siRNA and DICAR-JP detected by WB; D: OGDHL protein expressed in total cells; E: OGDHL protein expressed in mitochondria; F: OGDHL protein expressed in cytoplasm; G: Cell-free protein translation detecting the function of DICAR on OGDHL expression; H: DICAR-JP binding to OGDHL protein detected by RIP; I: SIM was used to detect the colocation of NACα, OGDHL, DICAR-JP with mitochondria, regulated by DICAR-JP. N = 3.

**Figure 3.**
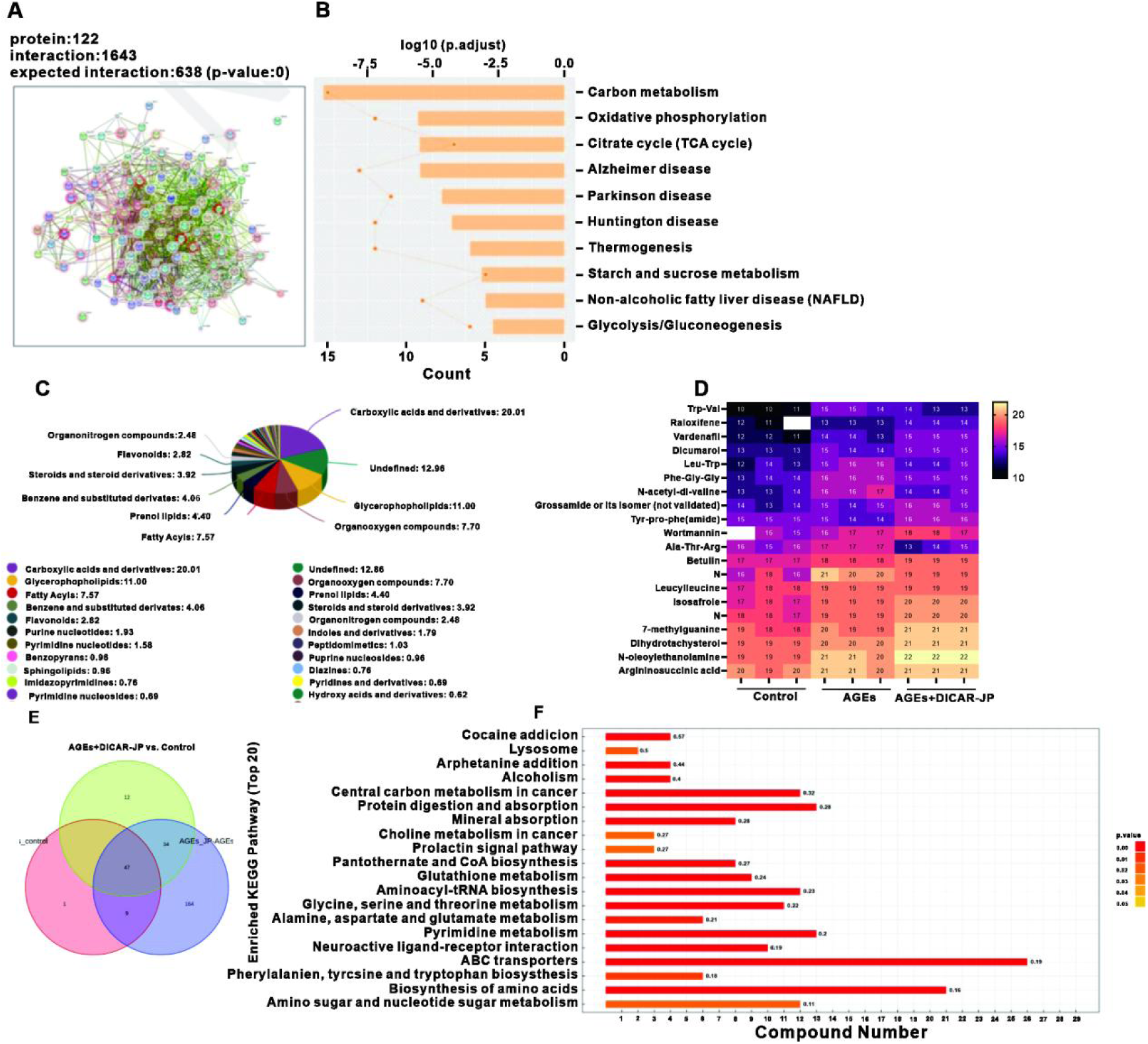
Effects of DICAR-JP on metabolic reprogramming induced by advanced glycation end products (AGEs) A: STRING ananlysed the interaction protein of energy metabolism binding with DICAR-JP; B: KEGG ananlyses the metabolism signal pathway; C: Pie chart ananlyses the classification of metabolism materials detected by untargeted metabolomics; D: Heat map showed the top ten changed metabolic materials in three groups; E: Venn diagram illustrating the altered metabolites across the three experimental groups; F: KEGG pathway analysis of the untargeted metabolomics data from AC16 cells treated with AGEs and DICAR-JP. N = 3.

### DICAR/DICAR-JP-OGDHL mediated cardiomyopathy metabolism reprogramming impaired with AGEs

As the KEGG analysis of DICAR CHIRP-MS, we found that the primary function of DICAR was significantly centered around its role in regulating various metabolic processes. It showed that there were 122 proteins playing interaction for eache other (Figure3A). Additionally, there were 10 metabolic signal pathway changed including carbon metabolism, oxidative phosphrylation, TCA cycle, et al (Figure 3B). To further confirmed the regulatory of DICAR-JP on metabolism impaired by AGEs, we employed untargeted metabolomics, to gain insights into the metabolic shifts associated with DICAR. The data of Figure3C demonstrated that AGEs contributing to metabolic dysregulation. As illustrated in Figure 3C, a total of 21 metabolic pathways were significantly altered with key pathways involving carboxylic acids and derivatives (accounting for 20.01% of the changes), glycerophospholipids (11.00%), organooxygen compounds (7.70%), fatty acyls (7.57%), and benzene and substituted derivatives (4.06%). Of the metabolites analyzed, 46 metabolic materials were found to be significantly altered across the three experimental groups, indicating substantial metabolic shifts (Figure 3D). Further pathway analysis using the KEGG revealed that the majority of these changes were linked to the ABC transport system, which plays a crucial role in the translocation of various substrates across membranes and comprised 26 altered members (Figure 3E). The second most affected pathway was the biosynthesis of amino acids, with 21 members showing significant changes. This suggests a broader implication of DICAR-JP’s role in fundamental metabolic processes, extending to amino acid synthesis and transport. Taken together, these findings suggest that DICAR-JP could be a promising therapeutic candidate, particularly in the context of diabetic cardiomyopathy (DCM), where metabolic dysregulation is a hallmark.

### DICAR-JP single base mutation is more suitable for binding with NACα

To confirm its DICAR-JP RNA sequence playing the protective function, not other RNA sequence, we selected different RNA sequence including DICAR junction site and synthesis them (Table S1). We tansfected DICAR-JP and DICAR-JPN(1-9) into AC16 to find the protective function on AGEs treated cell model detected by CCK8. The results showed that DICAR-JP increaed the cells abilitiy which was in accordance with our previous research. DICAR-JP3 and DICAR-JP9 exerted better function on cell activation (Figure 4A). Therefore, we chose DICAR-JP to optimize the activation, while DICAR-JP3 and DICAR-JP9 were left for another research. To explore potential improvements, we hypothesized that a single base mutation within the DICAR-JP sequence might enhance its interaction with NACα, a critical mediator in the regulatory pathway which can be used as a therapeutic function. We generated 66 RNA sequence variants of DICAR-JP and employed the HDOCK computational tool to predict and evaluate the binding affinities between DICAR-JP and NACα. Our analysis identified nine unique sequence variants of DICAR-JP, which we designated as DICAR-JPN (1-9) (Table 1, Figure 4B). These variants were individually transfected into AC16 human cardiomyocyte cells, and we evaluated cell viability using the CCK8 assay. The results revealed that exposure to AGEs (200 µg/ml) significantly reduced cellular activity, consistent with AGEs’ role in promoting oxidative stress and apoptosis. However, when AC16 cells were pretreated with the DICAR-JPN variants 24 hours prior to AGE exposure, only the DICAR-JP45 variant conferred a noticeable protective effect, enhancing cell viability compared to the control (Figure 4C). Unfortunately, at 48h, the protective effects of all DICAR-JPN variants, including DICAR-JP45, diminished (Figure 4C). Despite this, it demonstrated that DICAR-JP45 be the most promising initial results, prompting us to select it for further in-depth investigation in subsequent experiments. We tested the ED50 of DICAR-JP and DICAR-JP45, the results showed that the ED_50_ of DICAR-JP45 was 17.93 nM which was less than DICAR-JP EC50 26.19 nM (Figure 4D). DICAR-JP45 also belonged better effect on inhibiting pyroptosis of AC16 treated by AGEs (200 µg/ml 48 h, Figure 4E). We used SPR to find the interaction of DICAR-JP and DICAR-JP45. The results showed that DICAR-JP45 has better binding ability with NACα than DICAR-JP (Figure 4F). This study highlights the potential of DICAR-JP45 as a therapeutic nucleic drug candidate, though the diminishing protective effects underscores the need for further optimization of its sequence and delivery approach. Future work will focus on elucidating the mechanistic underpinnings of DICAR-JP45’s action and exploring additional modifications to prolong its protective benefits.

**Figure 4.**
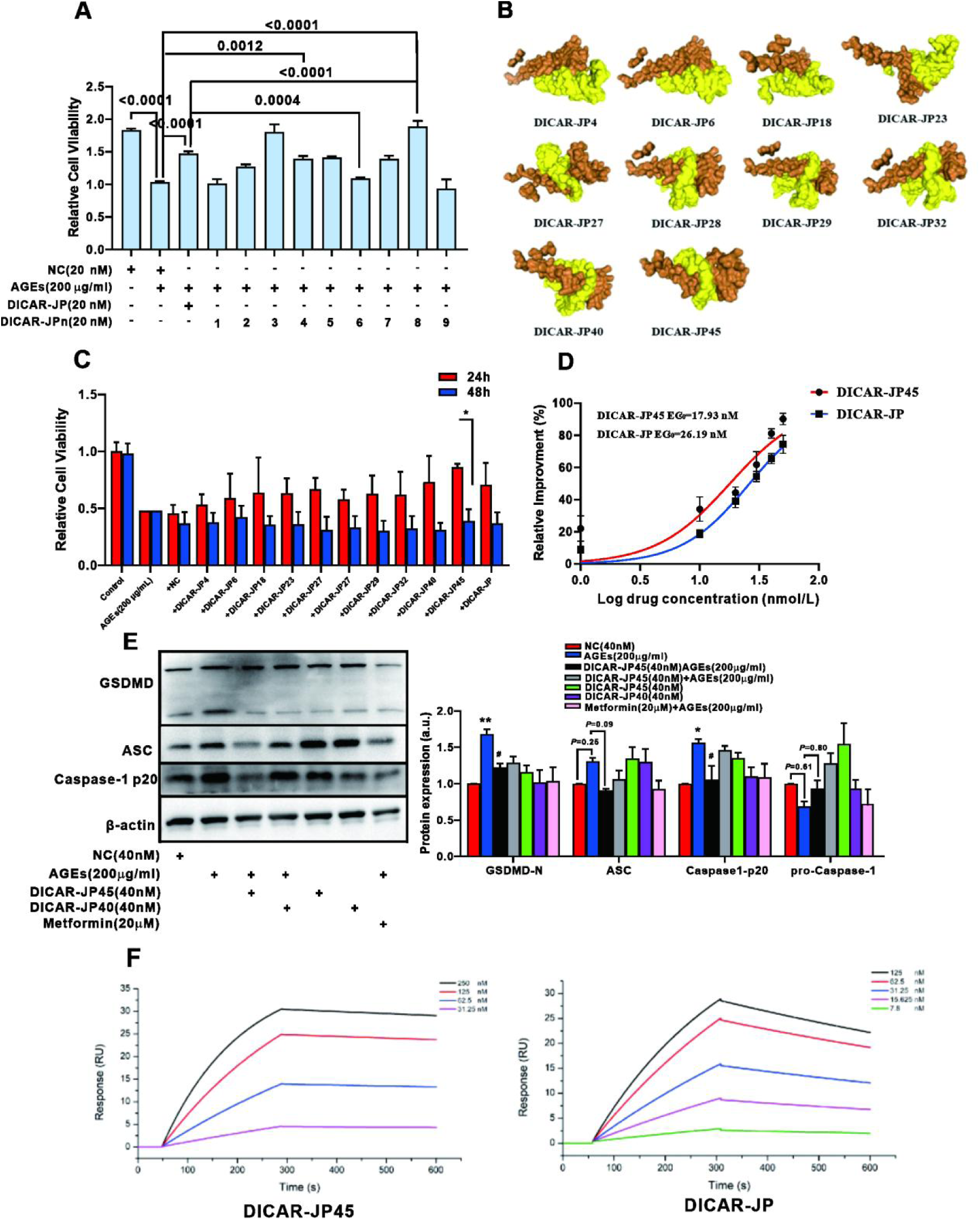
Optimization of DICAR-JP sequence with single base mutation. A: CCK8 detected DICAR-JP and DICAR-JPN on AC16 activation; B: Molecular binding interaction of DICAR-JPN with NACα; C: Protective effects of DICAR-JPN on AC16 cell viability as assessed by the CCK-8 assay; D: Dose-response curve depicting the EC50 values for DICAR-JP and DICAR-JP45; E: Representative images and statistical summary of pyroptosis induced by AGEs in AC16 cells; F: Synergistic effects of the combination of DICAR-JP and NACα. N = 3.

### AAV9-DICAR-JP45 played a beneficial effect on diabetic cardiomyopathy

We constructed two adeno-associated viral vectors, AAV9-DICAR-JP and AAV9-DICAR-JP45, and injected them into both db/m and db/db mice via tail vein injections to assess their therapeutic potential. Dapagliflozin (DAPA), a known SGLT2 inhibitor, was used as a positive control at a dosage of 1.5 mg/kg/day (Figure 5A). We successfully founded DICAR-JP and DICAR-JP45 overexpression in heart tissue (Figure 5B). Throughout the experiment, we continuously monitored the body weight of the db/db mice (Figure S1A); however, none of the treatment groups, including those receiving AAV9-DICAR-JP, AAV9-DICAR-JP45 or DAPA, demonstrated any significant effect on body weight regulation (Figure S1A). In terms of glucose metabolism, neither AAV9-DICAR-JP nor AAV9-DICAR-JP45 were able to significantly downregulate blood glucose levels in db/db mice, indicating that their therapeutic benefits might not be related to direct modulation of glycemic control (Figure S1C). We detected the heart function by Doppler. However, we did not observe the significantly heart dysfunction in db/db mouse model (data not shown), and therefore the protection of candidate drugs was not observed. Therefore, we considerated that heart function be not a good evalutaion indicator to db/db mouse aged 6 months. To further assess the structural and histological changes in the myocardium, wheat germ agglutinin (WGA) staining was performed to evaluate cardiomyocyte size, and Masson’s trichrome staining was used to detect myocardial fibrosis. Both AAV9-DICAR-JP and AAV9-DICAR-JP45 treatments attenuated pathological cardiac remodeling, including reducing cardiomyocyte hypertrophy and collagen deposition, indicating their effectiveness in mitigating diabetic cardiomyopathy (Figure 5D-E). These results suggest that AAV9-DICAR-JP45 played a more beneficial effect on heart function and remodeling compared with DAPA in DM. AAV9-DICAR-JP45 convey its protection effect by promoting OGDHL expression in the heart tissue (Figure 5E). DICAR-JP45 significantly inhibited NLRP3, ASC and cleaved-GSDMD expression of db/db heart tissue (Figure 5F).

**Figure 5.**
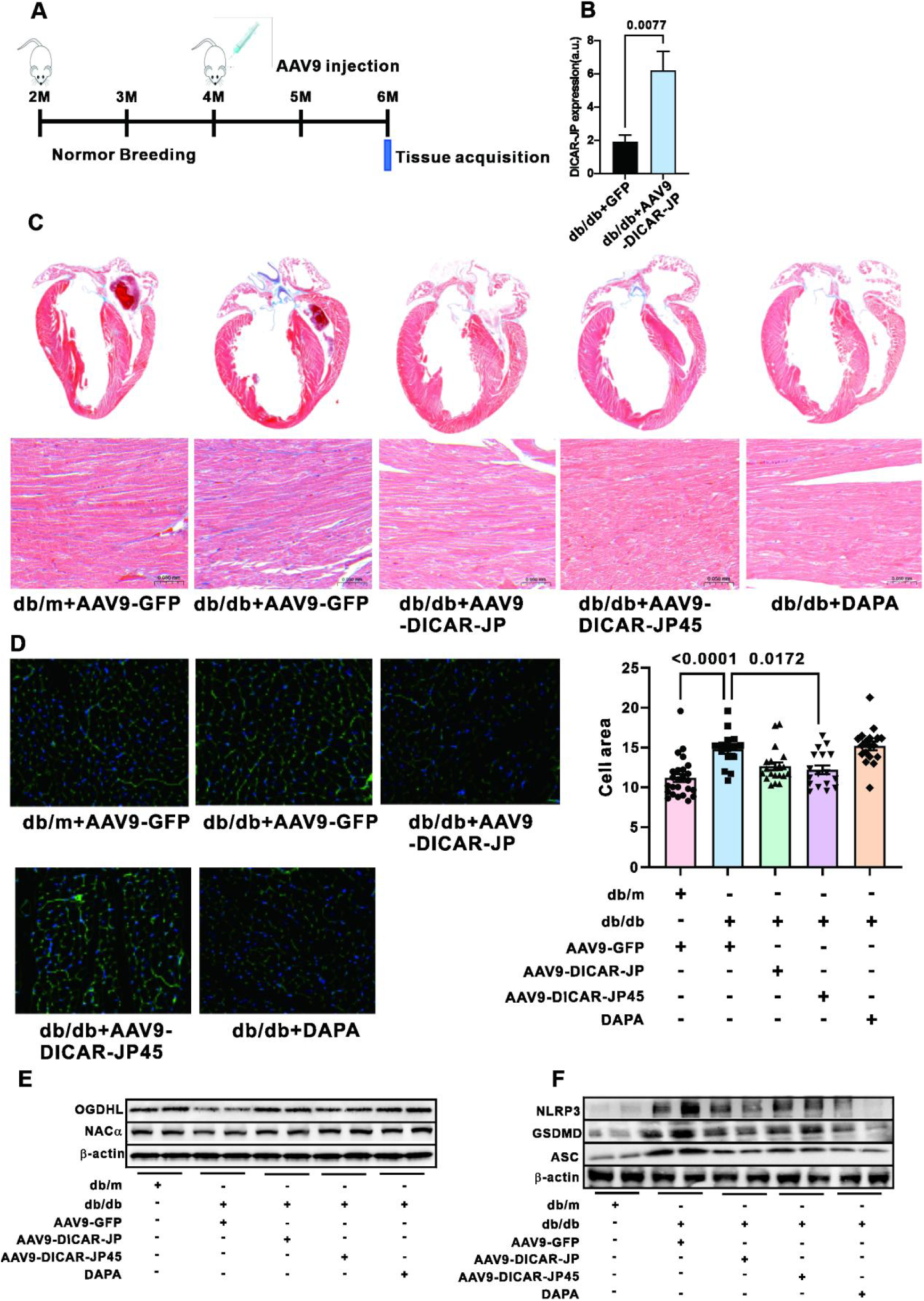
AAV9-DICAR-JP/JP45 protects cardiac function. A: Flowchart illustrating the treatment protocol for db/db mice with AAV9-DICAR-JP/JP45; B: Expression levels of DICAR-JP in cardiac tissue; C: Representative images of Masson stainning of cardiac tissue; D: Representative images of WGA stainning of cardiac tissue; E: OGDHL and NACα protein expression in heart tissue. N = 8.

## Discussion

This study suggests that DICAR-JP as the main functional domain of DICAR acts as a regulatory nucleic acid domain that binds to NACα, modulating its role in promoting the translation of OGDHL nascent polypeptides near the ribosomal exit tunnel. Furthermore, DICAR-JP facilitates the migration of ribosomes from the ER to the mitochondrial membrane, thereby enhancing the translation and translocation of OGDHL nascent polypeptides into mitochondria—a process we refer to as ‘ribosome migration’. Our results also demonstrate that the upregulation of OGDHL mediates mitochondrial energy dysregulation associated with diabetes. Based on the ‘functional nucleic acid’ hypothesis, we introduced a single-base modification in the DICAR-JP sequence, resulting in an optimized variant, DICAR-JP45, which exhibited enhanced efficacy as a potential nucleic acid-based therapeutic candidate for DCM.

NACα is a highly conserved alpha subunit of a heterodimeric complex called NAC. Recently, NACα has been reported to confer cellular susceptibility to reactive oxygen species (ROS) likely via negatively regulating the expression of the genes encoding Yap1, Skn7, Hog1, and Nox, all involved in ROS resistance^25^. At the ribosome exit site, NAC gates the activity of other nascent polypeptide chaperones. For example, NAC enhances the fidelity of SRP binding to only those nascent polypeptides destined for import to the ER. Therefore, all the functions of NACα are based on regulating nascent polypeptide production. In our research, we have identified that DICAR-JP binding with NACα regulates OGDHL nascent polypeptide production. Traditionally, all the researchers considered that ribosome are divided into two classes: ER ribosome and mitochondrion ribosome (55s). However, in this research, we found that DICAR-JP mediating NACα located in ER ribosome, and it migrated to the out membrane of mitochondria, promoting OGDHL translation and translocated into mitochondria, that is the translation and translocation steps are happened simultaneously. This protein was translated shared the same pathway of mitochondrial original proteins. We named this phenomenon ‘Ribosome Migration’, and identified another pathway to form mitochondrial proteins. Additionally, we propose a new concept ‘functional nucleic acid’. In our case, it means DICAR-JP is the effective functional nucleic part of circular RNA DICAR. Based on this theory, we eased the construction of circRNA with a shorter nucleic sequence. This helps with the discovery of nucleic drugs from ncRNAs for the next translational medicine research.

OGDHL is one of the rate-limiting components of the key mitochondrial multi-enzyme OGDH complex (OGDHC)^26^. Downregulation of OGDHL upregulated the α-ketoglutarate (αKG): citrate ratio by reducing OGDHC activity, which subsequently drove reductive carboxylation of glutamine-derived αKG via retrograde tricarboxylic acid cycling in hepatoma cells^26^. Additionally, OGDHL mRNA and protein displayed abnormal abundances in cardiac biopsies of HFpEF or heart failure with reduced ejection fraction patients^27,28–30^.

DICAR-JP functions as the junction site for circRNA-DICAR. In our previous investigations, we demonstrated that the transfection of DICAR-JP into AC16 cells significantly alleviated pyroptosis which was induced by advanced glycation end-products (AGEs). In vivo models, we postulated that the DICAR-JP sequence serves as a predicted binding site for NACα within its docking pocket. To facilitate this investigation, we designed specific primers for the detection of DICAR-JP expression in endogenous systems. Interestingly, our experimental analyses revealed the presence of DICAR-JP, suggesting that DICAR had undergone degradation to form this specific fragment. Furthermore, NACα was observed to exert a protective role over DICAR-JP, effectively inhibiting its degradation and thereby maintaining its functional integrity. Additionally, DICAR-JP functions as a structural molecular chaperone for NACα, regulating its activity and facilitating its translocation from the endoplasmic reticulum to the mitochondria. Moreover, we also elucidated an alternative pathway for the formation of ncRNAs, providing new insights into the complexities of RNA interactions and their regulatory mechanisms.

RNA therapies represent a significant advancement in the biomedical field, enabling the targeted manipulation of gene expression and the synthesis of therapeutic proteins. These innovative drugs are particularly well-suited for treating a range of pathologies characterized by established genetic targets, including infectious diseases, various malignancies, autoimmune disorders, and Mendelian genetic conditions^21^. The domain of nucleic acid therapeutics has emerged as a highly promising area of research within biotherapeutic methodologies. In recent years, several nucleic acid-based therapies have successfully transitioned to clinical application, especially for the treatment of metabolic and cardiovascular diseases. For example, Inclisiran has been shown to achieve a remarkable reduction of approximately 50% in low-density lipoprotein (LDL) cholesterol levels when administered subcutaneously every six months^22^. Similarly, Zilebesiran has been developed as a novel RNA interference therapeutic agent specifically targeting hypertension^23^. To further investigate the therapeutic potential of DICAR-JP, we introduced a targeted mutation at a single RNA base within the DICAR-JP sequence to enhance its binding affinity for NACα. Our experimental findings demonstrated that a single intravenous injection of DICAR-JP45, administered via the tail vein, conferred substantial protective effects on cardiac function in db/db mice over a duration of seven months. These protective effects were found to be comparable to those observed with both DICAR-JP and DAPA treatments. Moreover, DICAR-JP45 exhibited superior compliance relative to DAPA, with its beneficial effects maintained for nearly three months’ post-administration. Importantly, the protective efficacy of DICAR-JP45 was achieved without significant alterations in blood glucose levels or lipid metabolism, highlighting its potential as a safe and effective therapeutic agent. These findings underscore the promising role of DICAR-JP and its derivatives in the future development of RNA-based therapeutics, particularly in the context of cardiovascular health management.

In conclusion, this study proposed that the NACα/DICAR-JP complex plays a critical role in regulating the production of nascent polypeptides from OGDHL and facilitating ribosome migration. Furthermore, DICAR-JP is identified as a functional nucleic acid domain, while DICAR-JP45 emerges as a promising candidate for therapeutic nucleic acid applications in the treatment of diabetic cardiomyopathy (Figure 6).

**Figure 6.**
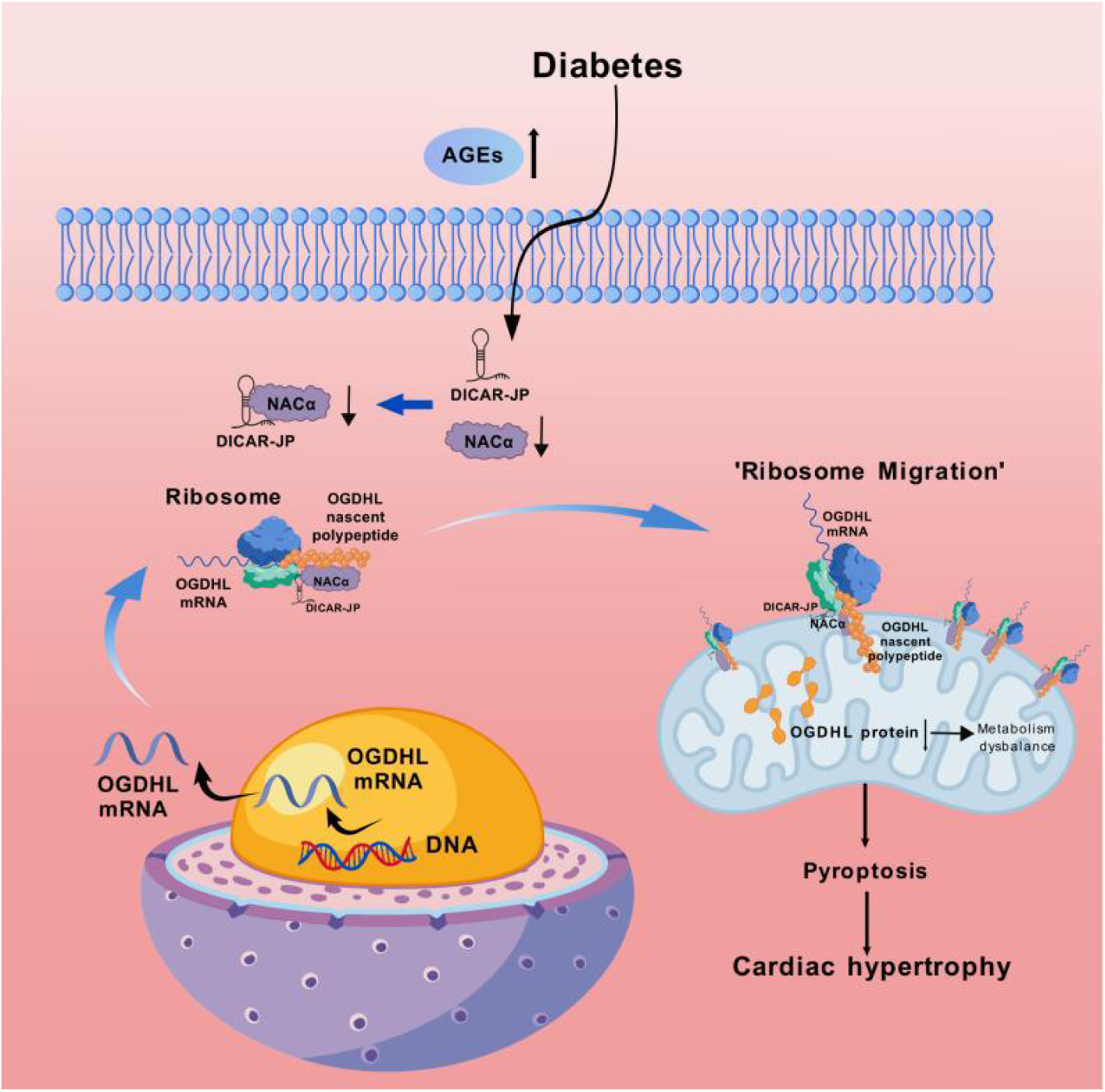
Proposed mechanism image.

## Nonstandard Abbreviations and Acronyms

DCM: diabetic cardiomyopathy
RIP: RNA immunoprecipitation
NACα: nascent polypeptide-associated complex α
OGDHL: oxoglutarate dehydrogenase-like
AGEs: advanced glycation end products
DAPA: dapagliflozin
SGLT2: sodium-glucose co-transporter-2
HFpEF: preserved ejection fraction
circRNA: circular RNA
VCP: valosin-containing protein
ER: endoplasmic reticulum
SRP: signal recognition particle
MS: mass spectrometry
FBS: fetal bovine serum
PMSF: phenylmethylsulfonylfluoride
SDS-PAGE: polyacrylamide gel electrophoresis
PVDF: polyvinylidene fluoride
ECL: chemiluminescent
CHIRP-MS: chromatin isolation by RNA purification–MS
KEGG: kyoto encyclopedia of genes and genomes
PCA: principal component analysis
OPLS-DA: orthogonal partial least squares-discriminant analysis
VIP: variable importance in projection
FC: fold change
TCA: tricarboxylic acid
IHC: immunohistochemistry
GSH: glutathione
LVVs: left ventricular volumes
WGA: wheat germ agglutinin
ROS: reactive oxygen species
LDL: low-density lipoprotein

## Acknowledgments

The authors thank Jie Gao for mouse heart function detected by Doppler. The authors thank SPR technical assistance of Wuhan Yangene Biological Technology Co, LTD, Yuechuang Center of HuaZhong Agricultural University.

## Author contributions

Conceptualization, Q.Y. and C.Z.; methodology, J.Z., X.Yu., C.Z., G.Q., X.G. and Q.Y.; investigation, S.T., Y.W., Y.Y., J.C., M.L., P.L., and Y.Y.; writing – original draft, Q.Y.; writing – review editing, C.Z., Q.Y. and Y.Y.; funding acquisition, Q.Y.; resources,C.Z., Q.Y.; supervision, C.Z., and Q.Y. J.Z., X.Yu., S.Z., G.Q., C.Z., S.T., X.G., Y.W., Y.Y., J.C., M.L., P.L., Y.Y., C.Z., Q.Y.

## Declaration of generative AI and AI-assisted technologies

During the preparation of this work, the author(s) used Chat-GPT in order to edit English language. After using this tool or service, the author(s) reviewed and edited the content as needed and takes full responsibility for the content of the publication.

